# Unveiling Ageing-Associated and Caloric Restriction-Associated Changes in miRNA Expression in Rat Skeletal Muscle and the Mechanisms Mediating their Effects on Muscle Cell Function

**DOI:** 10.1101/2024.04.07.588472

**Authors:** Gulum Altab, Brian J. Merry, Charles W. Beckett, Priyanka Raina, Ana Soriano-Arroquia, Bruce Zhang, Aphrodite Vasilaki, Katarzyna Goljanek-Whysall, João Pedro de Magalhães

## Abstract

The mechanisms underlying skeletal muscle ageing, whilst poorly understood, are thought to involve dysregulated micro (mi)RNA expression. Using young and aged rat skeletal muscle tissue, we applied high-throughput RNA sequencing to comprehensively study alterations in miRNA expression occurring with age, as well as the impact of caloric restriction (CR) on these changes. Furthermore, the function of the proteins targeted by these age- and CR-associated miRNAs was ascertained.

Numerous known and novel age-associated miRNAs were identified of which CR normalised 45.5% to youthful levels. Our results suggested miRNAs upregulated with age to downregulated proteins involved in muscle tissue development and metabolism, as well as longevity pathways, such as AMPK and autophagy. Furthermore, our results found miRNAs downregulated with age to upregulate pro-inflammatory proteins, particularly those involved in innate immunity and the complement and coagulation cascades. Interestingly, CR was particularly effective at normalising miRNAs upregulated with age, rescuing their associated protein coding genes but was less effective at rescuing anti-inflammatory miRNAs downregulated with age.

Lastly, the effects of a specific miRNA, miR-96-5p, identified by our analysis to be upregulated with age, were studied in culture C2C12 myoblasts. We demonstrated miR-96-5p to decrease cell viability and markers of mitochondrial biogenesis, myogenic differentiation and autophagy. Overall, our results provide useful information regarding how miRNA expression changes in skeletal muscle, as well as the consequences of these changes and how they are ameliorated by CR.

## Introduction

Skeletal muscle ageing is associated with the progressive deterioration of muscle mass, strength and function, also known as sarcopenia (Cruz-Jentoft et al. 2019). Ageing of skeletal muscle has a profound effect on the quality of life in older individuals. The loss of muscle mass begins approximately in the 4th decade of life and continues until the end of life. Studies indicate by the time an individual reaches the age of 80, there is a reduction of 30–50% in skeletal muscle mass and function (Akima et al. 2001). Human studies have demonstrated pension-age individuals to have 30– 40% less muscle strength and 25–30% less muscle size on average compared to young adults (Porter et al. 1995). The mechanisms that result in muscle ageing are not fully clear, and it is likely that numerous, interrelated processes drive the overall process. Some of the proposed mechanisms include reduced satellite cell number (Shefer et al. 2006), abnormal proteasomal degradation pathways (Fernando et al. 2019), increased ROS production (Palomero et al. 2013), mitochondrial dysfunction (Peterson et al. 2012), increased inflammation (Dalle et al. 2017) and abnormal expression long-non-coding RNAs (Marttila et al. 2020).

miRNAs are small non-coding RNAs, which negatively regulate gene expression post-translationally via mRNA repression and/or degradation. They have been identified as very important regulators in skeletal muscle proliferation and differentiation and have been demonstrated to play a role in age-related changes in skeletal muscle (Jung et al. 2017; Wang 2013; Drummond et al. 2011). Moreover, caloric restriction (CR), a widely studied method for combating aging in animals, seems to exert its influence on skeletal muscle, in part, by attenuating age-related changes in the miRNA pool (Margolis et al. 2016; Mercken et al. 2013). With this in mind, we sought to further investigate whether (1) miRNAs mediate muscle ageing, (2) CR can reverse age-associated miRNA pool changes in muscle and (3) pharmacological administration of select miRNA inhibitors/mimetics can reverse key markers of muscle ageing. Given, the human suffering caused by muscle ageing and subsequent sarcopenia, this knowledge could be of huge use with regards to improving the quality of life of older adults in future.

## Materials and methods

### Animals

Aged mice had a considerable reduction of muscular mass in the gastrocnemius and tibialis anterior (TA) muscles (Kim et al., 2014b). Thus, for this study particularly we investigated the gastrocnemius muscle. Gastrocnemius muscle tissue samples were collected from rats used in a previous study conducted by (Merry et al. 2008), under Home Office Project License 40/2964. All animal experiments were conducted in compliance with the United Kingdom Animals (Scientific Procedures) Act of 1986. In brief, 21-28 day old, male, Brown Norway (Substrain BN/SsNOlaHSD) rats were acquired from Harlan Laboratories, UK. Animals were kept under barrier conditions on a 12-h light:12-h dark cycle (08:00–20:00). Prior to 2 months of age, rats were group housed (n=4) and fed ad libitum (diet procured from Special Diet Systems Division Dietex International in Witham, UK). Following maturation, all animals were moved to solitary housing and randomly assigned to either ad libitum (AL) or caloric restriction (CR) diet groups. The CR rats were fed the same diet as the ad libitum rats, but their daily intake was restricted to 55% of the age-matched ad libitum intake. The CR rats were given access to food for a limited time each day between 10:30 and 11:00 hrs, while the ad libitum rats had unlimited access to the food for 24 hours per day. Animals were either terminated at 6 or 28 months of age (Table 1). No animals displayed any signs of clinical pathology prior to sacrifice. Following termination, gastrocnemius muscle tissue was removed, flash-frozen and stored at −80 °C, prior to further analysis.

**Table 1.**
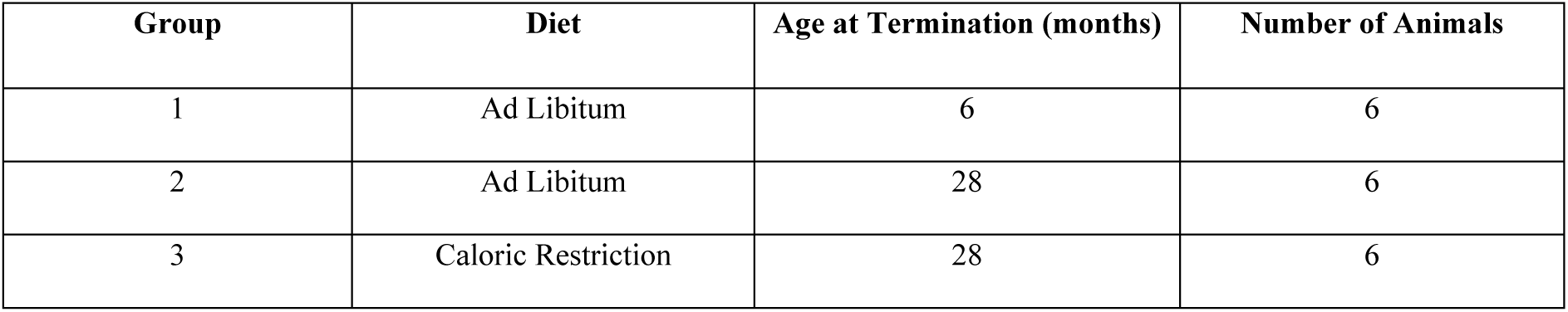
Group Identity, Diet Regimen and Age at Termination. Tissue samples were collected from male Brown Norway rats from one of 3 groups (n=6). Following animal arrival at 21-28 days of age, all animals were fed ad libitum (AL). After reaching full maturation/2-months old, animals were randomly allocated to either an AL (groups 1-2) or caloric restriction (CR) diet (group 3). Half the AL-fed animals were terminated at 6-months old and half at 28-months old. CR rats were terminated at 28-months old. Tissue samples were collected immediately post-sacrifice, snap frozen and stored at −80 °C until the day of analysis.

### RNA Extraction and RNA-Seq Analysis

Following removal from storage, gastrocnemius muscle tissues were immediately transferred onto ice to prevent thawing and degradation. Subsequently, each tissue was ground into a powder using a mortar and pestle. Liquid nitrogen was added periodically to ensure tissues remained in their frozen state. Total RNA was isolated using RNeasy Plus Universal Mini Kit (Qiagen, UK) according to manufacturer instructions. Extracted RNA quantity and quality were assessed on nanoDrop 2000 spectrophotometer and using the Agilent 2100 Bioanalyser, revealing all extracted RNA sampled to be of high quality (RINs>8). RNA integrity was further examined using agarose gel electrophoresis. Ribosomal RNA (rRNA) was removed from the total RNA using Illumina Ribo-Zero Plus rRNA depletion kit following the manufacturer’s instructions. At the Centre for Genomic Research (CGR), Liverpool, UK, utilizing the NEBNext small RNA library preparation Kit for Illumina, strand-specific RNASeq library preparation was performed. On the Illumina Hiseq 4000 platform, high-throughput RNA-seq was conducted (Paired-end, 2×150 bp sequencing, yielding data from >280 M clusters per lane). The raw Fastq files were trimmed for the presence of Illumina adapter sequences using Cutadapt version 1.2.1. The reads were further trimmed using Sickle version 1.200 with a minimum window quality score of 20. Sequencing and adapter trimming of data was conducted at CGR Liverpool, UK. Following trimming, shorter than 12-bp readings were removed.

The miRDeep2 method, which can identify known and unknown miRNAs in numerous independent samples with 98.6–99.9% confidence and accuracy (Friedlander et al., 2012), was used to identify novel and known miRNAs. The novel miRNAs were then filtered using the following two criteria: 1) miRDeep2 score >4. 2) significant randfold p-value (randfold refers to the likelihood that the predicted pre-miRNA sequence can fold into a stable RNA secondary structure). 3) At least two reads expressed in either young or old muscle samples. Although arbitrary, we chose a score of 4 to reduce the false positive detected miRNA.

### Differential Expression Analysis

Differentially expressed miRNAs between young AL-fed, old AL-fed and old CR rats were identified by using the Bioconductor package “DESeq2” (version, 3.7) in R (version 4.1.0). Raw gene counts, detected by featureCount, were used as the input. DESeq2 is a well-known method, which offers statistical procedures for identifying differential expression by using negative binomial linear models for individual genes and uses the Wald test for significance analysis. The P-values from the Wald test were then adjusted using the Benjamini and Hochberg’s method for controlling the false discovery rate (p-adj). Differentially expressed miRNAs were defined as those with a padj-value < =0.05 and fold change (FC)>=1.5.

### Micro RNA Target Identification

MultiMiR (version 1.20.0) in R was used to identify likely miRNA targets using information from several databases, including as miRanda, miRDB, PITA, TargetScan, MicroCosm, DIANA-microT, ElMMo, and PicTar (Ru et al., 2014). To narrow down miRNA targets identified by MultiMiR, only the top 40% of targets by score (an arbitrary cutoff) were classed as targets and any targets not found to be differentially expressed in our skeletal muscle ageing RNA-seq data were excluded. Though arbitrary, this tiered filtering focused our analysis toward the miRNA targets identified by MultiMiR with the greatest potential involvement in muscle ageing.

For novel miRNAs identified in this study, we applied a slightly different methodology as the MultiMiR package is limited to known miRNAs. We took advantage of the miRDB database (Wong & Wang 2015), to obtain predicted target genes for each of the novel miRNAs. While arbitrary, we chose a miRDB score cutoff of >60 to retain targets for further analysis. This moderate threshold balanced our exploratory goal of evaluating a subset of predicted targets for each miRNA while excluding lower-confidence targets with scores below 60. The resulting target gene lists were then directly used as input for gene ontology and KEGG pathway enrichment analysis to predict the potential functions of the novel miRNAs. We also used this target scan method to identify further unknown targets of miR-96-5p.

### Target Gene Ontology and Pathway Analyses

For the identified target genes, Gene ontology (GO) and KEGG pathways enrichment analysis was performed using ClusterProfiler (version 3.14.3), which enables visualization of functional profiles (Yu et al., 2012). Revigo (http://revigo.irb.hr) was used to remove redundant GO terms, clustering them based on semantic similarity. For this study, we performed GO biological process (GO-BP) analysis in the general level (GO level 3-8). Input gene sets were compared to the background (all differentially expressed genes according to RNAseq) using the hypergeometric test to obtain the significant GO terms and KEGG pathways (p<0.05) and the p-values corrected for false discovery rate (FDR). padj<=0.05 in multiple testing, q-values are also projected for FDR control. The top 20 GO and KEGG pathways were then visualised using ggplot2 (version 3.3.5) in R.

### RT-qPCR Validation of miRNAseq Data Analysis

Once RNA-seq results were obtained regarding which genes were differentially expressed, we examined 11 of these differentially expressed miRNAs using quantitative real-time PCR (RT-qPCR) on the gastrocnemius muscles of young (6-month-old), old (24-month-old) and old CR (28-month-old) rat to validate the age-related differentially expressed miRNA discovered by miRNA sequencing (miRNAseq). This was done to validate the results of our RNA-seq analysis.

Validated Primers for miRNAs were obtained from QIAGEN:

| **Assay Name** | **Catalogue Number** |
| --- | --- |
| mmu-miR-31-5p | YP00205159 |
| hsa-miR-96-5p | YPO0204417 |

| Assay Name | Catalogue Number |
| --- | --- |
| rno-miR-380-3p | YPO2102544 |
| hsa-miR-101-3p | YPO0204786 |
| hsa-miR-122-5p | YP00205664 |
| hsa-miR-20a-5p | YP00204292 |
| hsa-miR-375 | YP00204362 |
| rno-miR-3576 | YP02103725 |
| hsa-miR-140-3p | YP00204304 |
| mmu-miR-182-5p | YP00205089 |

We amalgamated normalized data from samples of both young and old, as well as old CR, into a unified file. Subsequently, NormFinder (http://moma.dk/normfinder-software) was employed for analysis. Notably, mir-101a-3p emerged as one of the most consistently stable miRNAs across all samples and served as the internal control in our study..For RT-qPCR samples were calculated relative to rno-miR-101a-3p miRNA levels and group comparisons were conducted using One-Way ANOVA test followed by Newman-Keuls Multiple Comparison Test. The significance level was set to 5%.

### C2C12 Cell Culture and Transfection

We used culture myoblast populations to investigate the role of a miRNA (rno-miR-96-5p), revealed by our RNA-seq/bioinformatic analyses to be important to both muscle ageing and responses to CR. C2C12 cells were grown in high gluose Dulbecco’s Modified Eagle Medium (DMEM) supplemented with 10% foetal bovine serum (FBS) (Thermos Scientific, UK), 1% l-glutamine, and 1% penicillin/streptomycin. Once 90% confluent, the cells were grown in DMEM supplemented with 2% horse serum, 1% l-glutamine, and 1% penicillin/streptomycin, stimulating myogenic differentiation. Subsequently, using Lipofectamine 2000 (Thermos Scientific, UK), in triplicates C2C12 cells were transfected with either 50 nM of miR-96-5p, 50nM of a mock mimetic (control), 100 nM of a miR-96-5p inhibitor or 100nM of a scrambled inhibitor (control) (Qiagen, UK).

### Assessment of the Effect of miR-96-5p on Cell Viability and Myogenic Differentiation in C2C12 cells

Cell viability and differentiation were assessed in separate cell populations cultured and transfected using the method described above. After a 24h post-transfection incubation period, cell viability was assessed using acridine orange (Sigma-Aldrich, A8097) /ethidium bromide (Sigma-Aldrich, E1510) (AO/EB) staining (Kasibhatla et al. 2006). In brief, the cell media was removed and washed twice with PBS before being stained with 500ul of a PBS solution containing 1:1000 ethidium bromide and 1:1000 acridine orange. 3 images were obtained per condition.Images of the cells were acquired using a fluorescence microscope with a total magnification of 100x, utilising the green channel for acridine orange and the red channel for ethidium bromide. Following fluorescent microscopy, the total numbers of live and dead cells were determined using ImageJ (Schneider et al. 2012). Subsequently, the ratio of live to dead cells was calculated to assess the overall viability of each cell population.

In other cell populations, after a 7-day period post-incubation, MF20 (myosin heavy chain) immunostaining was conducted to quantify the level of myoblast differentiation. In brief, cells were stained with both DAPI (Sigma-Aldrich)and MF20 antibodies (MF20-c 2ea, DSHB). 3 images were obtained per condition (Soriano-Arroquia et al. 2017). Following fluorescent microscopy, quantitative measures of myoblast differentiation, including fusion index, MyHC positive area and myotube diameter, were derived. For Fusion index, number of nuclei within MF20 positive cells (cells containing more than 2 nuclei) were counted manually, and the total nuclei was counted using image J “Analyse Particles” automated function. Fusion index was calculated by dividing the number of nuclei within multinucleated myotubes by the total number of nuclei in the entire field of view. Meanwhile, for each treatment condition, Image J also facilitated the measurement of the MyHC positive areas by adjusting threshold, this gave the measurement of the total area covered by MyHC-positive staining relative to the total area of the field of view. The MYHC staining threshold in ImageJ was set by accessing the ‘Image’ menu, selecting ‘Adjust’, and then ‘Threshold’. Parameters were carefully adjusted to enhance MYHC positive staining visibility while minimizing background noise, ensuring effective isolation and highlighting of positive areas while mitigating the impact of extraneous background elements in the images. To measure the myotube diameter, firstly, the image containing myotubes was opened in ImageJ. A line selection tool was utilised to draw a straight line across the diameter of individual myotubes. The ‘Analyse’ menu was accessed, and ‘Set Scale’ was used to calibrate the image with known measurements. Subsequently, the ‘Straight Line’ tool was employed to measure the distance across the myotubes, and the recorded values were used to determine the myotube diameter.

### Assessment of the effect of miR-96-5p on OGDH and Mitochondrial Biogenesis Marker Expression in C2C12 cells

We hypothesised miR-96-5p to directly downregulate OGDH expression and mitochondrial biogenesis downstream. Therefore, using qPCR, we investigated the expression of OGDH, two key factors involved in mitochondrial biogenesis, peroxisome proliferator-activated receptor gamma coactivator 1-alpha (PGC-1α) and mitochondrial transcription factor A (TFAM), and cytochrome c oxidase subunit I (COX I), a key mitochondrial protein. The miRCURY LNA RT Kit (Qiagen, UK) was used to synthesise cDNA using 50-100ng of RNA, and the miRCURY LNA RT Kit (Qiagen, UK) was used to conduct qPCR in the Roche ® LightCycler® 480 according to the manufacturer’s instructions. Validated primers were obtained from Qiagen, UK for the miRNAs. rno-miR-101a-5p or U6 was used as an internal control. As for the protein coding genes, the following primers were used; OGDH F: GGTGTCGTCAATCAGCCTGAGT R: ATCCAGCCAGTGCTTGATGTGC (Origene, UK); PGC1α F: GAACAAGACTATTGAGCGAACC, R: GAGTGGCTGCCTTGGGTA; COX1 F: ACTATACTACTACTAACAGACCG, R: GGTTCTTTTTTTCCGGGAGT; 18S F: AGAAACGGCTACCACATCCA, R: CCCTCCAATGGATCCTCGTT (internal control) (Sharma et al., 2021); TFAM, F: GAGGCAAAGGATGATTCGGCTC, R: CGAATCCTATCATCTTTAGCAAGC (Fukaya et al., 2021).

### Assessment of the effect of miR-96-5p on Mitochondrial Function Using Citrate Synthase Assay

Proteins were extracted from C2C12 cells that were either treated with a control, miR-96-5p mimic or inhibitor for the purpose of conducting a citrate synthase assay. The MitoCheck® Citrate Synthase Activity Assay Kit from Cayman Scientific (USA) was utilized to measure the activity of citrate synthase. In the case of the ATP assay, C2C12 cells were transfected with control, miR-96-5p mimic, and inhibitors for a duration of 24 hours.

### Assessment of the effect of miR-96-5p on Autophagy Marker (LC3A, LC3B) Expression in C2C12 cells using Western Blotting

C2C12 protein was extracted from cells using RIPA buffer (Sigma-Aldrich, UK. Cat # R0278) using the manufacturer’s instructions. Following extraction, the isolated proteins were stored at −80°C prior to analysis. Subsequently, the BCA assay was used to determine total protein concentration. Firstly, the protein standard was placed on ice while a portion of the sample was diluted to 1:10. Then, the BSA standard curve was produced using the manufacturer’s recommendation. A working reagent of BCA solution was generated by mixing 50 parts of BCA Reagent A with 1 part of BCA Reagent B (50:1 ratio). In a 96-well plate (CoStar, USA), 20ul of protein standard and 20ul of sample proteins were added. Then, 180μl of BCA working reagent was added to each well and the plate incubated at 37°C in a dark room for 30 minutes. Following the incubation period, the plate was scanned using Spectro Star Nano (BMG LabTech, Germany) at a wavelength of 562 nm. The average absorbance was measured and utilised to quantify the protein concentration. Subsequently, SDS-PAGE electrophoresis was performed.

Following preparation of the polyacrylamide gels, it was carefully placed into the electrophoresis tank. Subsequently, cell protein samples and a protein ladder were loaded into their respective wells within the gel. The tank was filled with 1X Tris-Glycine SDS PAGE Buffer diluted in distilled water to facilitate the electrophoretic process. Additionally, protein loading buffer (prepared by mixing tracking dye at a 1:6 ratio) was added to the running buffer, aiding in the visualization of sample migration. The electrophoresis began at a lower voltage, typically set around 60V initially, to allow the samples to traverse the stacking gel. Once the bromophenol dye migrated to approximately 2mm from the bottom of the gel, indicating separation had occurred, the voltage was increased to 80V. At this point, the running of the gel was halted.

For Western blotting, buffers were prepared for the transfer of proteins onto nitrocellulose membranes using the semi-dry transfer technique. A current was applied for transfer, followed by verification with Ponceau S staining. The membrane underwent blocking with 5% skim milk for 1 hr, followed by immunoblotting utilising LC3 antibodies (cell signalling technology, cat: 2775) and multiple washes before analysis using the LICOR Cx system.

### Dual-Luciferase Reporter and ATP assays

Target scan using the https://mirdb.org/custom.html database (Wong and Wang, 2015) revealed significant alignment between the 3’UTR segment of OGDH and miR-96-5p in several species (Figure 19). Consequently, to confirm the interaction between miR-96-5p and OGDH, we created a luciferase reporter pmirGLO vector containing a segment either the OGDH 3’UTR, containing the purported miR-96-5p binding site (the OGDH wild-type; WT), or a mutant version of the OGDH 3’UTR devoid of the miR-96-5p binding site (OGDH Mutant; MUT). C2C12 cells were co-transfected with scRNA control or miR-96-5p mimic, together with the pmirGLO vector containing either WT or MUT OGDH 3’UTR.

Wild type (“CTTTGCTGTGCCAAGGCA”) and mutated (“CTTTGCTTTGCCGGGGCA”) seed sequence 3’UTR of OGDH were used (Thermos Scientific, UK). RT-PCR was used to amplify the OGDH 3’UTR and sub clone it into pmirGLO vector (Promega, UK). 5 × 10^4 C2C12 cells were seeded into 12 well plates. The next day, using Lipofectamine 2000, cells were co-transfected with control or miR-96-5p mimic and pmirGLO luciferase vector carrying either the OGDH 3’UTR wild-type or mutant gene. Dual-luciferase reporter gene assays were performed 24 hours post-transfection according to the protocol used in a previous study (Yang et al., 2015). The level of ATP under different conditions was determined using the Mitochondrial ToxGlo™ Assay from Promega (UK), following the protocol provided by the manufacturer.

### Statistical Analysis of Cell Culture Experiments

RT-qPCR data were analysed using the delta-delta Ct technique (2–Ct method) to determine relative expression levels between treatment conditions (Rao et al. 2013). The mean ± SEM of the results were plotted using Prism (5.01 edition) for Windows. To examine the difference between the groups, an unpaired two-tailed Student’s t-test was conducted.. For the cell viability study, an unpaired two-tailed Student’s t-test was conducted to compare the average live:dead cell ratio across treatment conditions. For the myogenic differentiation study, one-way analysis of variance, followed by Newman-Keuls Multiple Comparison test, were conducted to compare average fusion index, myotubule diameter and MyHC positive area between treatment conditions.

Likewise, all additional statistical analyses were performed using Prism (5.01 edition), with a significance level established at 5%. In the case of Western blot, citrate synthase assay, and luciferase assay, an Unpaired two-tailed t-test was employed to assess differences between the control and treatment groups.

## Results

### Identification of novel miRNAs expressed in rat skeletal muscle and their targets

We analysed skeletal muscle of young (6-month-old) and old (28-month-old) AL-fed rats, and old CR (28-month-old CR) rat to examine the ageing process. We obtained small RNA reads derived from 18 miRNA libraries, multiplexed, and sequenced over 2 lanes of the HiSeq 4000 (2×150), generating an average of ∼31M paired end reads. Following trimming, miRDeep2 identified 146 novel miRNAs in rat skeletal muscle, all of which, given their significant randfold value, folded into the typical hairpin shape of a mature miRNA. The miRDeep2 algorithm found that several novel miRNAs shared homology in their seed sequence with existing miRNAs found in mice, suggesting a likely similarity. MultiMiR proposed numerous protein-coding genes to be the targets of the identified miRNAs.

### Identification of miRNAs differentially expressed between the muscle of young and aged AL-fed rats

We identified a total of 84 miRNAs that were significantly differentially expressed (FDR corrected padj-value ≤0.05, FC 1.5) in the muscle of aged rats fed AL compared to young rats fed AL. 36 miRNAs were significantly upregulated, of which 31 miRNAs were known and 5 were newly discovered by miRDeep2. Meanwhile, 48 were significantly downregulated, of which 41 were known and 8 were newly discovered by miRDeep2. The top miRNAs exhibiting the greatest degree of upregulation (Table 2A) included rno-novel-132, rno-novel-79, rno-novel-78, and rno-novel-101 (novel) and rno-miR-31a-5p, rno-miR-122b, rno-miR-122-5p, rno-miR-122-3p, rno-miR-182, rno-miR-146a-5p and rno-miR-9b-5p (known). The top miRNAs exhibiting the greatest degree of downregulation (Table 2B) included rno-novel-70, rno-novel-84, rno-novel-88 and rno-novel-39 (novel) and rno-miR-539-5p, rno-miR-3576, rno-miR-369-5p, rno-miR-3578, rno-miR-434-3p, rno-miR-3543, rno-miR-376b-5p, rno-miR-136-5p, rno-miR-181d-5p, rno-miR-410-3p and rno-miR-181c-5p (known).

**Table 2.** Top 20 (A) upregulated and top 20 (B) downregulated miRNAs in old rat skeletal muscle compared with young rat skeletal muscle, ranked by padj-value. Rats were split into two groups (n=6/group), which were terminated at either 6-months or 28-months old. Gastrocnemius muscle tissue was removed from rats terminated at either 6-months or 28-months old and RNA was extracted and sequenced. DESeq2 in R, analysing raw gene counts via negative binomial models and Wald test, identified differentially expressed miRNAs among young AL-fed and old CR rats, considering significance via adjusted p-values (padj) ≤ 0.05 and fold change (FC) ≥ 1.5 criteria. Differentially expressed miRNAs were defined as those with a padj-value < =0.05 and fold change (FC)>=1.5.

|  |  |  |  |  |
| --- | --- | --- | --- | --- |
| (A) | miRNA | log2FoldChange | padj | BaseMean |
|  | rno-novel-132 | 25.95507919 | 6.34E-16 | 357.1308 |
|  | rno-novel-79 | 24.75568433 | 1.29E-14 | 151.9206 |
|  | rno-novel-78 | 22.93445537 | 1.42E-12 | 41.09418 |
|  | rno-novel-101 | 21.52469818 | 4.19E-11 | 14.89469 |
|  | rno-miR-31a-5p | 3.600712113 | 1.40E-08 | 24.79392 |
|  | rno-miR-122b | 4.473785927 | 6.26E-06 | 3592.289 |
|  | rno-miR-122-5p | 4.330237239 | 1.11E-05 | 7951.759 |
|  | rno-miR-598-3p | 1.195398731 | 0.000116064 | 175.9525 |
|  | rno-miR-140-3p | 0.653852793 | 0.00023907 | 18190.15 |
|  | rno-miR-122-3p | 4.259059829 | 0.000329713 | 52.70041 |
|  | rno-miR-182 | 1.162119257 | 0.000577317 | 1379.57 |
|  | rno-miR-375-3p | 2.695336409 | 0.000723014 | 34.71595 |
|  | rno-novel-47 | 10.86271201 | 0.000837844 | 190.9947 |
|  | rno-miR-483-3p | 2.654748309 | 0.001388372 | 380.344 |
|  | rno-miR-212-3p | 1.950123623 | 0.001446286 | 9.452453 |
|  | rno-miR-129-1-3p | 2.506946968 | 0.001446286 | 11.65899 |
|  | rno-miR-146a-5p | 0.747539925 | 0.004345804 | 15795.15 |
|  | rno-miR-193b-3p | 1.250428873 | 0.004507445 | 309.1669 |
|  | rno-miR-9b-5p | 1.879357786 | 0.008134343 | 14.10007 |
| (B) | miRNA | log2FoldChange | padj | BaseMean |
|  | rno-novel-70 | -30 | 5.86E-21 | 108074.1 |
|  | rno-novel-84 | -30 | 5.86E-21 | 11186.94 |
|  | rno-novel-88 | -29.68172627 | 1.13E-20 | 3814.042 |
|  | rno-novel-39 | -26.67203183 | 1.15E-16 | 290.9535 |
|  | rno-novel-52 | -26.1424763 | 4.41E-16 | 198.9091 |
|  | rno-novel-104 | -25.72144436 | 1.07E-15 | 370.7493 |
|  | rno-novel-105 | -25.29484489 | 3.20E-15 | 105.9255 |
|  | rno-novel-113 | -23.56296473 | 3.04E-13 | 29.80431 |
|  | rno-novel-109 | -22.89962192 | 1.46E-12 | 18.23865 |
|  | rno-miR-539-5p | -2.253351594 | 4.38E-10 | 29.70962 |
|  | rno-miR-3576 | -2.253842252 | 1.14E-09 | 29.70669 |
|  | rno-miR-369-5p | -1.492813542 | 6.22E-09 | 85.70875 |
|  | rno-miR-3578 | -1.491103379 | 6.22E-09 | 86.08394 |
|  | rno-miR-434-3p | -1.232290361 | 4.70E-06 | 1138.526 |
|  | rno-miR-3543 | -1.231824278 | 5.43E-06 | 1131.442 |
|  | rno-miR-376b-5p | -1.944052377 | 2.17E-05 | 40.65792 |
|  | rno-miR-136-5p | -1.730578207 | 6.21E-05 | 83.56321 |
|  | rno-miR-181d-5p | -1.087475984 | 0.000329713 | 818.2401 |
|  | rno-miR-410-3p | -1.337094704 | 0.000414971 | 55.57651 |

### Identification of miRNAs differentially expressed between the muscle of aged CR and aged AL-fed rats

We identified a total of 109 miRNAs that were significantly differentially expressed (FDR corrected padj-value ≤0.05, FC 1.5) in the muscle of aged rats fed CR compared to young rats fed AL. 49 miRNAs were significantly upregulated, of which 41 miRNAs were known and 8 were newly discovered by miRDeep2. Meanwhile, 60 were significantly downregulated, of which 53 were known and 7 were newly discovered by miRDeep2. The top 20 miRNAs exhibiting the greatest degree of upregulation (Table 3) and downregulation (Table 4) were ascertained.

**Table 3.** Top 20 upregulated miRNAs in old calorically restricted (CR) rat skeletal muscle compared with young ad libitum (AL)-fed rats, ranked by padj-value. Rats were split into two groups (n=6/group) and fed either ad libitum (AL) and terminated at 6-months old or CR and terminated at 28-months old. Gastrocnemius muscle tissue was removed from which RNA was extracted and sequenced. DESeq2 in R, analysing raw gene counts via negative binomial models and Wald test, identified differentially expressed miRNAs among young AL-fed and old CR rats, considering significance via adjusted p-values (padj) ≤ 0.05 and fold change (FC) ≥ 1.5 criteria. Differentially expressed miRNAs were defined as those with a padj-value < =0.05 and fold change (FC)>=1.5.

| miRNA | log2FoldChange | padj | BaseMean |
| --- | --- | --- | --- |
| rno-novel-100 | 26.64661238 | 1.60E-16 | 1345.95 |
| rno-novel-145 | 24.68883073 | 2.28E-14 | 136.8081 |
| rno-novel-66 | 14.99782935 | 8.12E-13 | 3186.408 |
| rno-novel-78 | 22.76976921 | 2.59E-12 | 34.60824 |
| rno-novel-102 | 21.76249613 | 2.47E-11 | 16.82333 |
| rno-miR-31a-5p | 4.670883357 | 2.94E-09 | 47.75662 |
| rno-miR-598-3p | 0.995735699 | 3.69E-09 | 151.6259 |
| rno-miR-346 | 3.294970618 | 8.79E-09 | 36.07669 |
| rno-novel-12 | 0.892388925 | 2.38E-06 | 6764.193 |
| rno-miR-139-5p | 0.88927553 | 2.81E-06 | 6881.724 |
| rno-miR-129-5p | 2.632092541 | 3.49E-06 | 126.2047 |
| rno-miR-9b-5p | 3.106562825 | 5.44E-06 | 27.43186 |
| rno-miR-122b | 3.661126884 | 1.82E-05 | 1986.573 |
| rno-novel-49 | 6.483718798 | 1.82E-05 | 8.726301 |
| rno-miR-122-5p | 3.542309879 | 2.81E-05 | 4482.465 |
| rno-miR-9b-3p | 2.29252938 | 3.01E-05 | 1973.445 |
| rno-miR-9a-5p | 2.286957031 | 3.02E-05 | 2033.489 |
| rno-miR-129-1-3p | 3.648988731 | 3.54E-05 | 18.51977 |
| rno-miR-9a-3p | 2.724083389 | 4.19E-05 | 214.8999 |

**Table 4.** Top 20 downregulated miRNAs in old calorically restricted (CR) rat skeletal muscle compared with young ad libitum (AL)-fed rats, ranked by padj-value. Rats were split into two groups (n=6/group) and fed either ad libitum (AL) and terminated at 6-months old or CR and terminated at 28-months old. Gastrocnemius muscle tissue was removed from which RNA was extracted and sequenced. DESeq2 in R, analysing raw gene counts via negative binomial models and Wald test, identified differentially expressed miRNAs among young AL-fed and old CR rats, considering significance via adjusted p-values (padj) ≤ 0.05 and fold change (FC) ≥ 1.5 criteria. Differentially expressed miRNAs were defined as those with a padj-value < =0.05 and fold change (FC)>=1.5.

| miRNA | log2FoldChange | padj | BaseMean |
| --- | --- | --- | --- |
| rno-novel-88 | -30 | 1.14E-20 | 3640.95 |
| rno-novel-104 | -26.81122264 | 1.47E-16 | 352.7158 |
| rno-novel-81 | -26.36557812 | 2.78E-16 | 254.3544 |
| rno-novel-105 | -25.07195011 | 9.38E-15 | 99.09345 |
| rno-novel-113 | -23.38232807 | 6.81E-13 | 28.58696 |
| rno-novel-109 | -22.71193274 | 2.75E-12 | 17.4121 |
| rno-miR-181c-3p | -1.386804517 | 9.06E-12 | 51.03027 |
| rno-miR-493-5p | -1.443916102 | 1.5E-10 | 121.4076 |
| rno-miR-1193-3p | -1.587389832 | 3.08E-09 | 46.19807 |
| rno-miR-539-5p | -1.824437777 | 0.000000042 | 29.84787 |
| rno-miR-3576 | -1.811380362 | 0.000000109 | 29.90049 |
| rno-miR-181d-5p | -1.066464749 | 0.000000119 | 782.6765 |
| rno-miR-434-3p | -1.38068115 | 0.000000141 | 1049.362 |
| rno-miR-3543 | -1.379247764 | 0.000000145 | 1043.056 |
| rno-miR-337-5p | -1.266920566 | 0.000000225 | 309.3306 |
| rno-miR-181c-5p | -1.763366261 | 0.000000326 | 82.40578 |
| rno-miR-329-3p | -1.529038698 | 0.000000377 | 105.5133 |
| rno-miR-409a-5p | -1.231325898 | 0.0000129 | 102.6213 |
| rno-miR-471-3p | -5.33316902 | 0.0000226 | 3.481599 |

### Age-related miRNA changes in old skeletal muscle of rats partially reversed by caloric restriction

Of the 84 miRNAs differentially expressed between aged rats fed AL and young rats fed AL, 34 (40.5%) were normalised to youthful expression levels in aged CR rats. Of the 36 miRNAs upregulated in the muscle of aged rats fed AL compared to young rats fed AL, 17 (47.2%) were suppressed in aged CR rats. Therefore, CR failed to suppress 19 (52.8%) of the miRNAs upregulated with ageing. Of the 48 miRNAs downregulated in the muscle of aged rats fed AL compared to young rats fed AL, 17 (35.4%) were rescued in aged CR rats. Therefore, CR failed to rescue 31 (64.6%) of the miRNAs downregulated with ageing.

**Figure 1.**
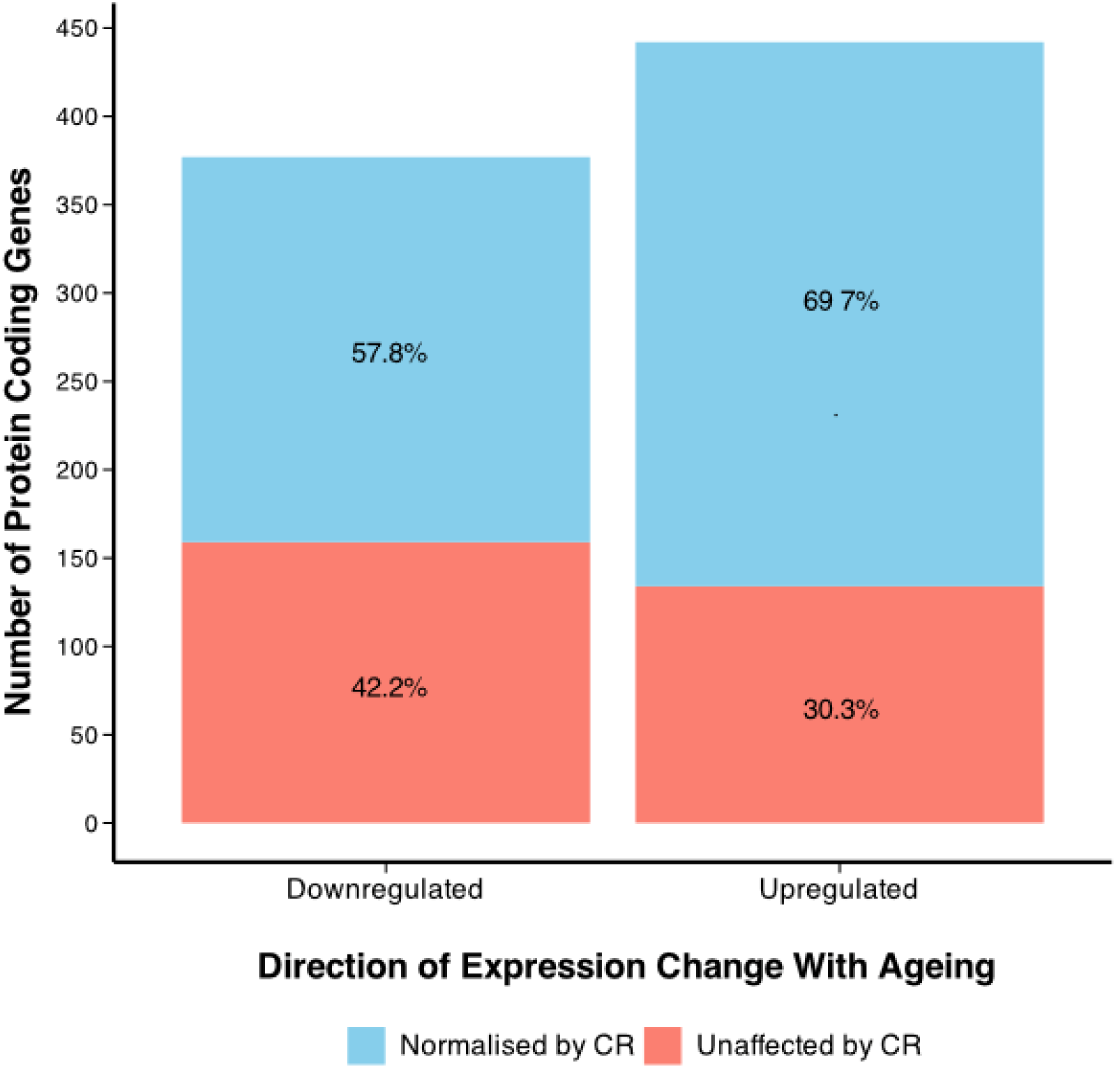
Impact of CR on gene expression. A bar chart showing the number of genes upregulated and downregulated in the muscle of aged rats fed ad libitum (AL), when compared to the muscle of old rats fed AL, and what percentage of these are normalised by caloric restriction (CR). Venn diagrams describe the number of genes. POTENTIALLY ADD TWO TABLES TO SUPPLEMENTARY MATERIAL

### Functional and pathway analysis of protein-coding genes targeted by miRNA differentially expressed in aged rat muscle

Known upregulated miRNAs in old skeletal muscle targeted 175 protein-coding genes, which were enriched for 39 GO terms. Notable terms included cellular response to peptide hormone stimulus, skeletal muscle development, glycogen metabolic process, and skeletal muscle cell differentiation. REVIGO streamlined redundant terms, summarising target genes to be involved in skeletal muscle development, positive regulation of fatty acid oxidation, and fructose/hexose metabolism (Figure 3A). Lastly, KEGG pathway analysis revealed miRNAs to target genes associated with cGMP-PKG signalling pathway, AMPK signalling pathway, Longevity regulating pathway, Insulin resistance, Adipocytokine signalling pathway and Autophagy signalling pathways (Figure 3B).

**Figure 3.**
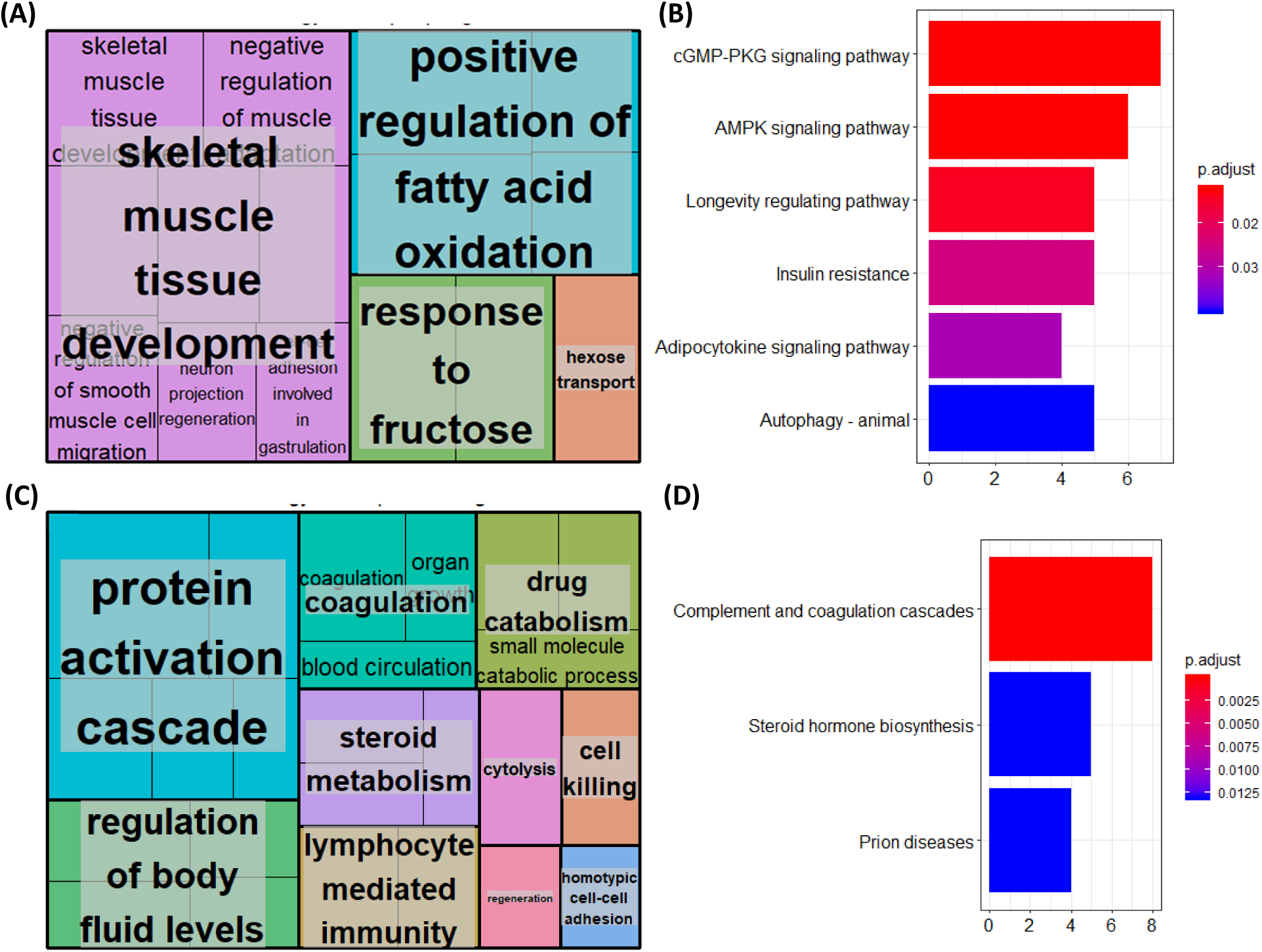
Functions and pathways of genes targeted by known miRNAs upregulated (A-B) and downregulated (C-D) in the muscle of aged rats compared to young rats. Gastrocnemius muscle from 6-months or 28-months old rats was isolated and RNA extracted and sequenced. Differentially expressed miRNAs were revealed using Desq2. For the known target genes of differentially expressed miRNAs, gene ontology (GO) terms, which were clustered based on semantic similarity (A, C) and KEGG pathway enrichment were derived.

Known downregulated miRNAs in old skeletal muscle targeted 202 protein-coding genes, which were enriched for 60 GO terms. Notable terms included wound healing, response to molecules of bacterial origin, response to lipopolysaccharide, regulation of body fluid levels and lymphocyte-mediated immunity. REVIGO streamlined redundant terms, summarising target genes to be involved in the protein activation cascade, regulation of body fluids, coagulation, drug metabolism, steroid metabolism and lymphocyte-mediated immunity, amongst others (Figure 3C) Lastly, KEGG pathway analysis revealed miRNAs to target genes associated with complement and coagulation cascades, steroid biosynthesis and prion diseases (Figure 3D).

Analysis of novel miRNAs upregulated in aged rat muscle using the miRDB database revealed numerous target genes. These were found to be significantly associated with six GO terms: axonogenesis, stress-activated MAPK cascade, regulation of BMP signalling pathway, and glial cell migration (Figure 4A). KEGG pathway analysis highlighted miRNAs to potentially target of crucial skeletal muscle pathways, including PI3K-Akt, MAPK, and mTOR signalling (Figure 4B).

**Figure 4.**
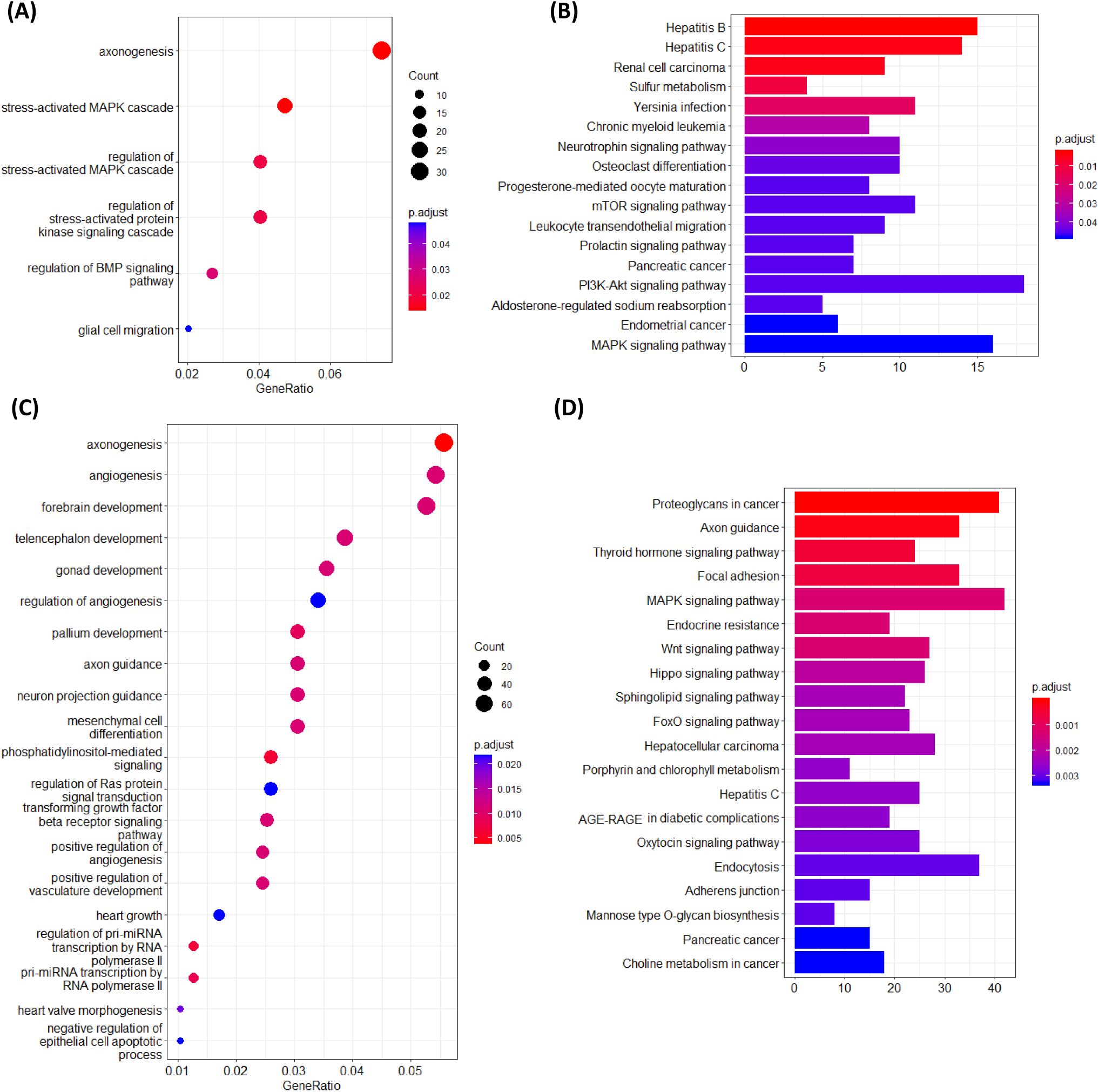
Functions and pathways of genes targeted by novel miRNAs upregulated (A-B) and downregulated (C-D) in the muscle of aged rats compared to young rats. Gastrocnemius muscle tissue was removed from rats terminated at either 6-months or 28-months old and RNA extracted and sequenced. Differentially expressed miRNAs were revealed using Desq2. Novel miRNAs were analyzed using the mirdb.org database to predict target genes based on a cutoff score >60, facilitating subsequent gene ontology (A, C), and KEGG pathway enrichment (B, D) analyses to infer potential functions.

Analysis of novel miRNAs upregulated in aged rat muscle using the miRDB database revealed numerous target genes. These were found to be significantly associated with 43 GO terms, notably including axonogenesis, angiogenesis, and forebrain development (Figure 4C). KEGG pathway analysis for downregulated miRNAs identified 61 significantly associated pathways, notably including Proteoglycans in cancer, Axon guidance, MAPK signalling, Cellular senescence, and several other pathways (Figure 4D).

### Functional and pathway analysis of protein-coding genes targeted by miRNA uniquely differentially expressed in the muscle of aged CR rats compared to young AL-fed rats

The miRNAs uniquely upregulated by CR were enriched for 13 GO terms. Notable terms included positive regulation of cellular carbohydrate metabolic process, positive regulation of glucose metabolic process, regulation of polysaccharide biosynthetic process and regulation of glucose metabolic process. REVIGO summarised these as related to carbohydrate metabolism and regulation of lamellipodium organisation (Figure 5A). KEGG pathway enrichment analysis was not performed due to the low quantity of resultant GO terms.

**Figure 5.**
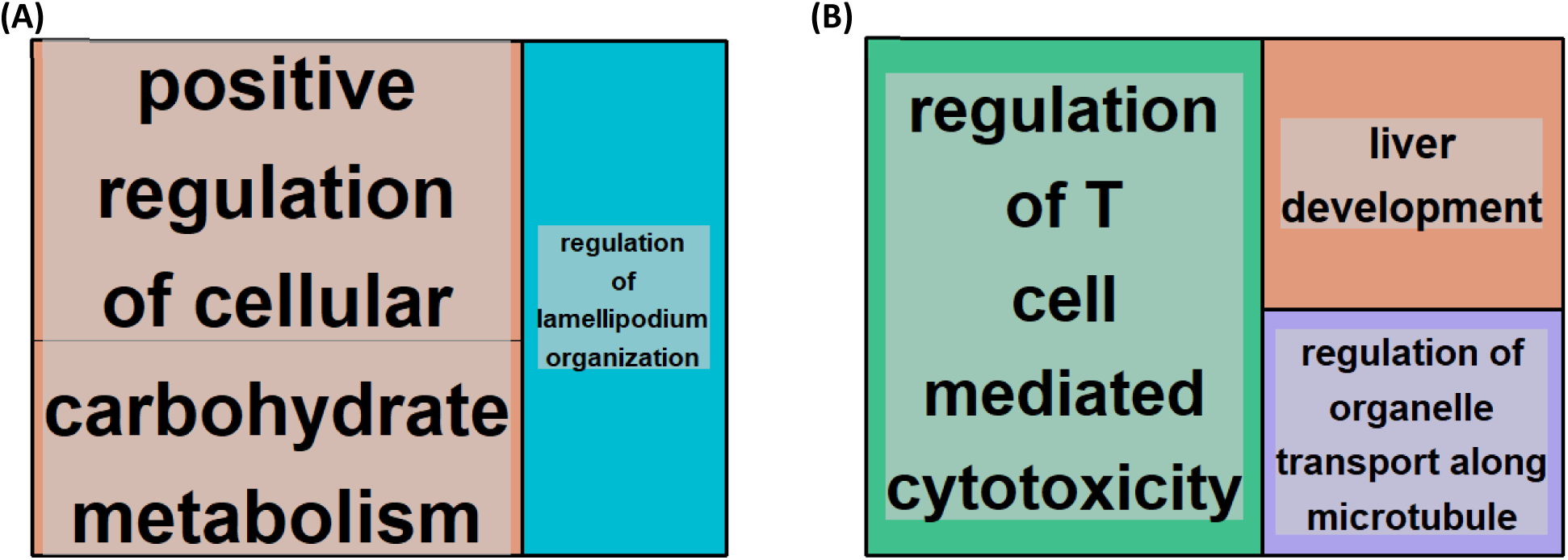
Gene ontology (GO) terms associated with genes targeted by miRNAs uniquely upregulated (A) and downregulated (B) in the muscle of aged calorically restricted CR rats compared to young ad libitum (AL)-fed rats. Gastrocnemius muscle tissue was removed from rats either fed AL and terminated at 6-months or CR and terminated at 28-months old. RNA was extracted and sequenced. Differentially expressed miRNAs were revealed using Desq2 and target genes determined and associated GO terms, which were clustered based on semantic similarity (A, C).

The miRNAs uniquely downregulated by CR were enriched for 9 GO terms. REVIGO summarised these as involved in regulation of T cell-mediated cytotoxicity, liver development and regulation of organelle transport along microtubules (Figure 5B). Lastly, KEGG pathway analysis revealed miRNAs to target genes associated with Allograft rejection, Graft-versus-host disease, Autoimmune thyroid disease, Type I diabetes mellitus, Cellular senescence, Viral myocarditis, Antigen processing and presentation, Epstein-Barr virus infection, Cell adhesion molecules (CAMs) amongst other pathways (Figure 6).

**Figure 6.**
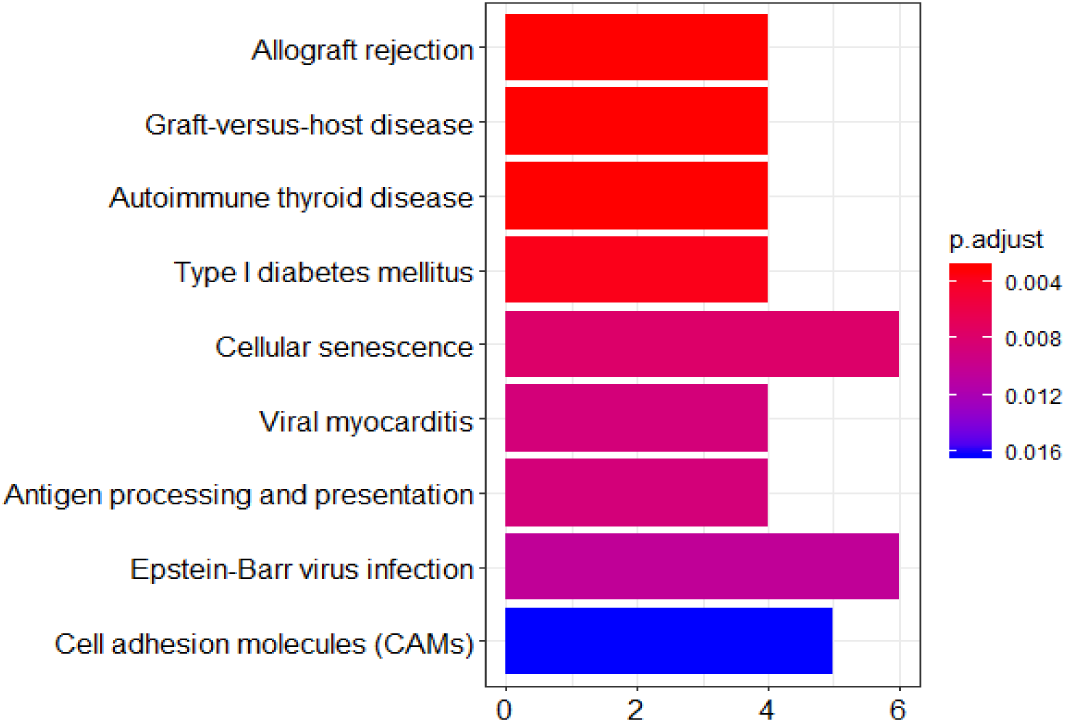
KEGG pathways associated with genes targeted by miRNAs uniquely downregulated in the muscle of aged calorically restricted CR rats compared to young ad libitum (AL) rats. Gastrocnemius muscle tissue was removed from rats and RNA extracted and sequenced. KEGG pathways associated with differentially expressed miRNAs were revealed using Desq2 and ClusterProfiler.

### RT-qPCR Validation of miRNAseq data analysis

We validated differential expression of 7 miRNAs, rno-miR-122-5p, rno-miR-96-5p, rno-miR-375-5p, rno-miR-20a-5p, rno-miR-31a-5p, rno-miR-140-3p, and rno-miR-182, using qPCR. Four out of seven of the miRNAs showed identical results with miRNAseq data analysis. These included rno-miR-122-5p, rno-miR-96-5p, rno-miR-31a-5p and rno-miR-140-3p. These miRNAs were upregulated in old rat skeletal muscle compared to young rat skeletal muscle, and expression of these miRNAs was inhibited by CR (Figure 7 A-D, I-L).

**Figure 7.**
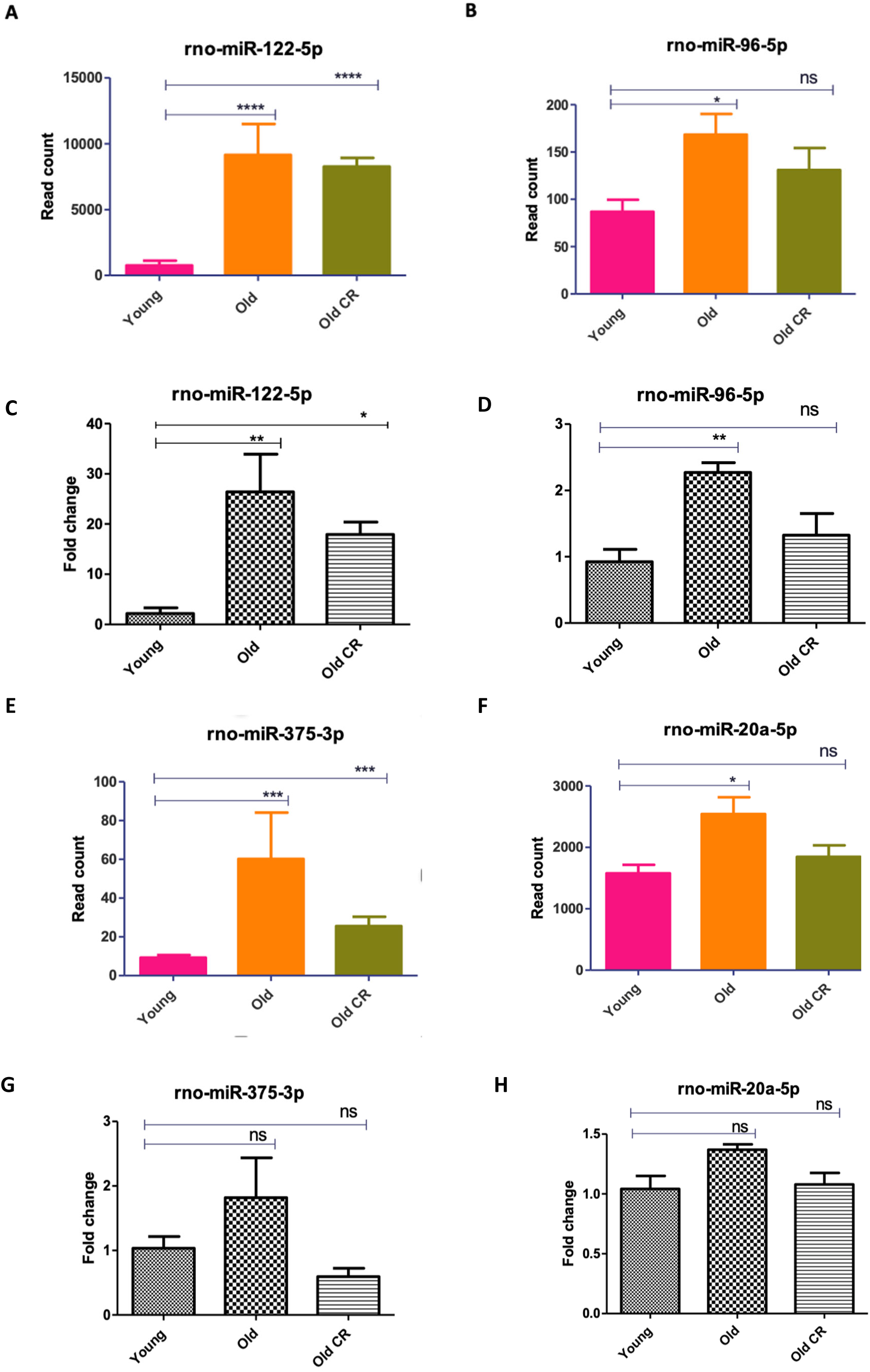
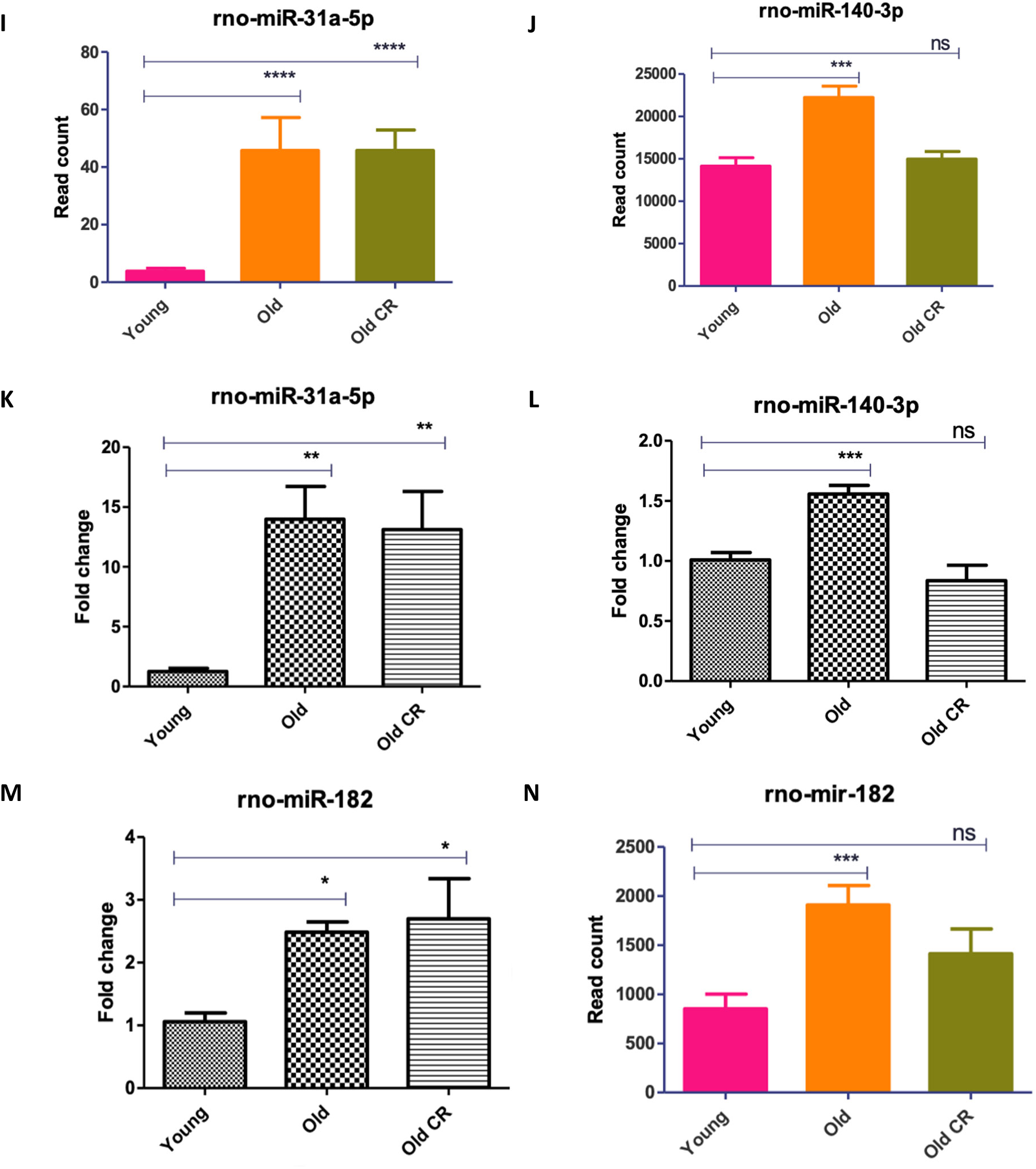
RT-qPCR validation of miRNA-seq results. RNA was extracted from the gastrocnemius muscle of all rats and analysed using both RNA-seq and qPCR. For RT-qPCR samples differential expression was calculated in fold change relative to a housekeeper miRNA, rno-miR-101a-5p. Graphs showing normalised read count from microRNA-seq data. Group comparisons were conducted using One-Way ANOVA test followed by Newman-Keuls Multiple Comparison Test. (A-B,E-F,I-J, M) show miRNA-seq results and (C-D,G-H,K-L, N) show RT-qPCR results. All p-values are unadjusted for rt-qpcr. Error bars represent mean ± SEM, n = 4–6, ns denotes not significant,*=0.01 to 0.05, **= 0.001 to 0.01, ***= 0.0001 to 0.001 and ****= <0.0001.

Whilst RT-qPCR results showed the remainder of miRNAs to exhibit similar expression trends in response to ageing and CR, they did not exactly replicate RNA-seq results. For example, RT-qPCR found rno-miR-375-5p and rno-miR-20a-5p to not be significantly differentially expressed between animal groups (Figure 7 E-H). Furthermore, whilst RT-qPCR found rno-miR-182 (Figure 7 M,N) to be upregulated in the muscle of aged rats fed AL (replicating RNA-seq results), no significant difference was found between expression in young rats fed AL and old CR rats (contrary to RNA-seq results). Nevertheless, overall RT-qPCR analysis showed consistent results with miRNAseq differential analysis, thus, demonstrating the reliability of our finding.

### Rationale for further investigating rno-miR-96-5p

The miRNA rno-miR-96-5p has previously been shown to inhibit myogenic differentiation via suppression of Four-and-a-half LIM domains protein 1 (FHL1) gene expression (10.3390/ijms21249445). Interestingly, our RNA-seq and bioinformatic analyses highlighted differential expression of this miRNA to likely be a significant player both in the muscle aging process and in the response to CR. Its target genes were found to be involved in several aspects of optimal skeletal muscle function, such as responses to insulin, promotion of fatty acid oxidation and tissue development. Furthermore, its target genes were associated with a number of longevity-associated functions, including AMPK signalling, autophagy and responses to advanced glycation end products (AGEs). Being upregulated in aged rat muscle, rno-miR-96-5p likely plays a role in skeletal muscle dysfunction with age via downregulating many of these longevity-associated genes. We, therefore, sought to further investigate the effects of rno-miR-96-5p on myoblast function by administering a miR-96-5p mimetic and inhibitor to cultured C2C12 cells.

### The effect of miR-96-5p on viability and myogenic differentiation in cultured C2C12 cells

AO/EB staining revealed there to be no significant effect of 50nM miR-96-5p on the ratio of live to dead cells compared to the population receiving 50nM of the mock mimetic. However, the cell population transfected with 100nM of the miR-96-5p inhibitor had a significantly higher ratio of live to dead cells compared to the population receiving 100nM of the mock inhibitor (Figure 8A).

**Figure 8.**
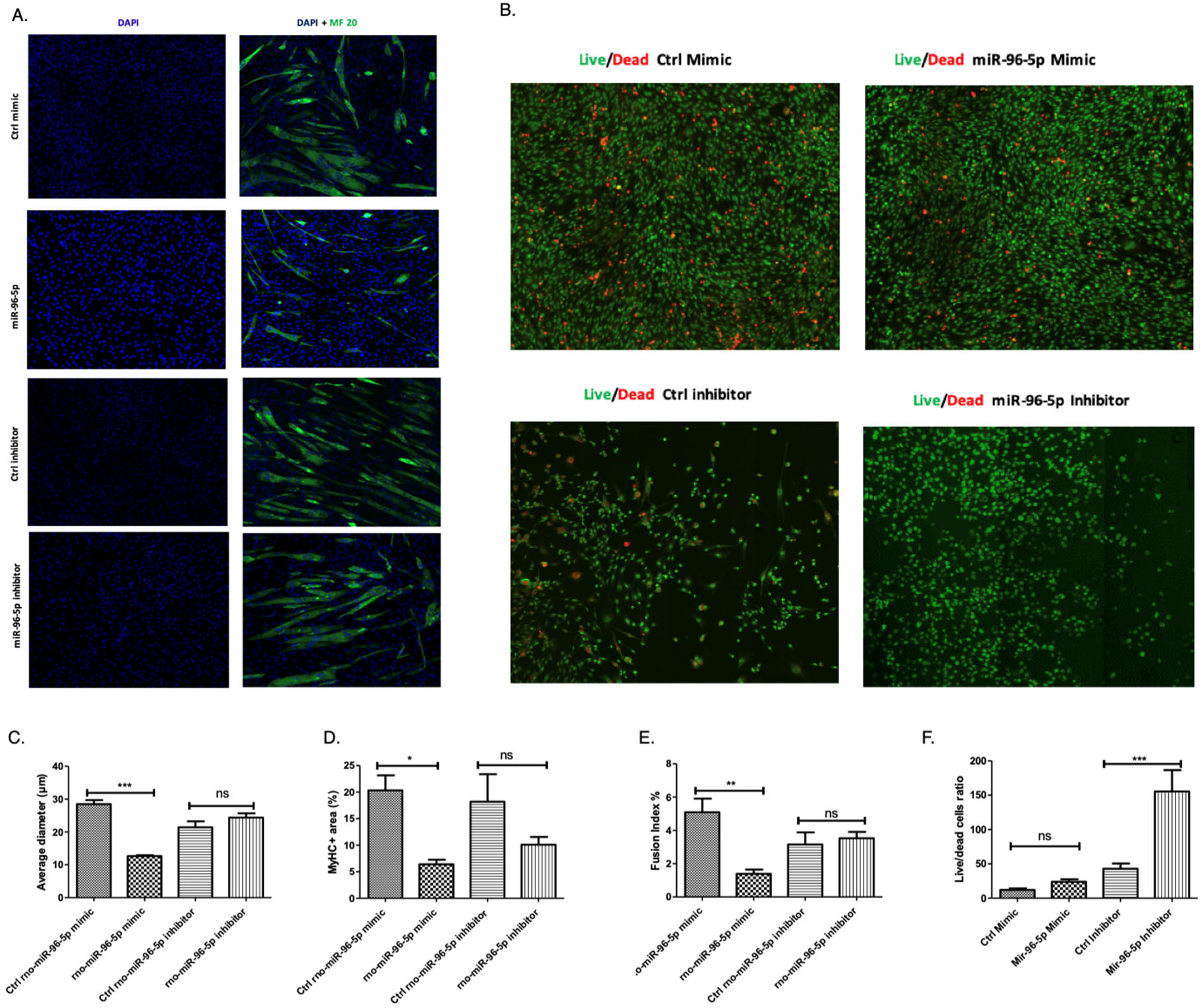
Myogenic differentiation and cell viability are reduced by over-expression of miR-96-5p in C2C12 cells. C2C12 myoblasts were transfected with either 50nM of a control mimetic, 50nM of miR-96-5p mimetic, 100nM of a control inhibitor or 100nM of a miR-96-5p inhibitor. (A, C-E) Myoblast nuclei were stained with DAPI and heavy chain of myosin II was stained with an immunofluorescent MF20 antibody. Myotube diameter (C) and MyHC positive area (D) were ascertained using ImageJ. The fusion index (E) was quantified by dividing the number of multinucleated myotubes by the total number of nuclei in the entire field of view across multiple images and analysed by one-way analysis of variance and Newman-Keuls Multiple comparisons test. (B, F) Acridine orange/ethidium bromide revealed the number of live and dead cells in each cell population. The average live:dead cells ratio was calculated for each treatment condition and compared using unpaired two-tailed Student’s t-tests. All statistical tests were conducted in Prism. Error bars represent mean ± SEM, n = 3-4, ns denotes not significant, *=0.01 to 0.05, **= 0.001 to 0.01, ***= 0.0001 to 0.001 and ****= p-value <0.0001.

MF20 (myosin heavy chain) immunostaining revealed the cell population transfected with 50nM miR-96-5p to have a significantly lower fusion index, myotubule diameter and MyHC positive area compared to the cell population transfected with 50nM of the mock mimetic indicating this miR negatively regulates myogenesis. No other significant group effects were found for any measure. However, there was a non-significant trend toward a lower MyHC positive area in cells receiving the miR-96-5p inhibitor (Figure 8B-D). It is possible that the effects of inhibition of miR-96 may need to be explored in stress conditions to uncover a potential functional effect.

### The effect of miR-96-5p on mitochondrial biogenesis markers

qPCR revealed that cells transfected with miR-96-5p to have no significantly lower expression of TFAM compared to control transfected cells. No significant differences between these groups were observed with regard to the expression of PGC-1α or COX I. Cells transfected with the miR-96-5p inhibitor displayed significantly higher expression levels of PGC-1α, COX I and TFAM compared to cells receiving the mock inhibitor (Figure 9A).

**Figure 9.**
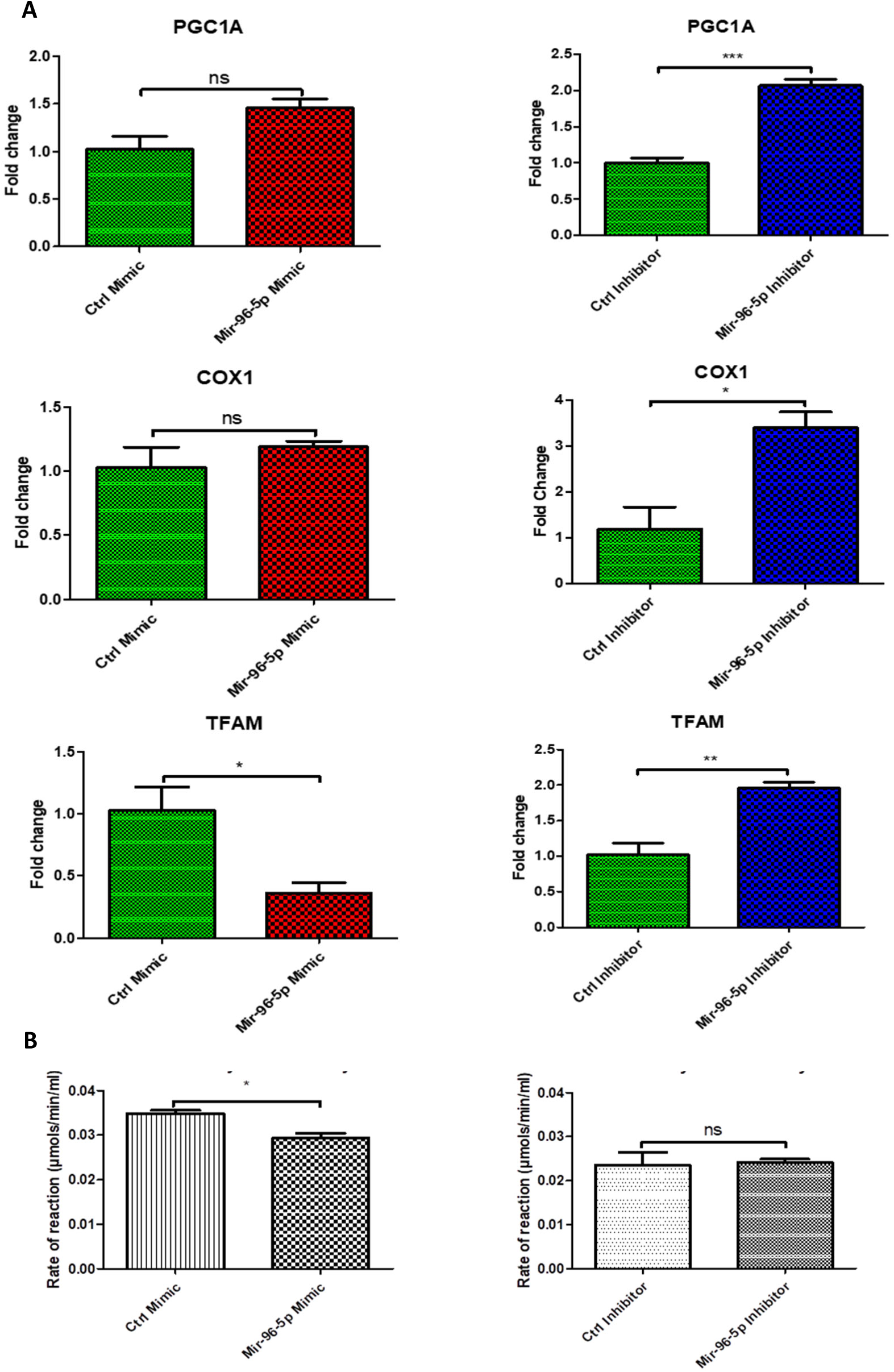
Effect of miR-96-5p on markers of mitochondrial biogenesis. C2C12 myoblasts were transfected with 50 nM of control mimic, 50nM miR-96-5p mimic, 100nM control inhibitor or 100nM of a miR-96-5p inhibitor. (A) RT-qPCR was conducted 24h after transfection. Expression of PGC-1α, COX1 and TFAM in C2C12 was quantified relative to 18S. (B) A citrate synthase assay was conducted using MitoCheck. The average activity for each treatment condition was determined in μmols/min/ml. Error bars represent mean ± SEM, n = 3,*=p-value<0.01 to 0.05, **= 0.001 to 0.01, ***= 0.0001 to 0.001.

To confirm our findings, we performed a citrate synthase activity assay, which is used to determine the activity of mitochondrial citrate synthase in samples. This revealed that cells transfected with miR-96-5p to have significantly lower citrate synthase activity compared to cells transfected with the control mimetic. However, no significant difference was observed between cells treated with the control inhibitor versus the miR-96-5p inhibitor (Figure 9B).

### The effect of miR-96-5p on autophagy markers

Western blot results revealed both LC3A and LC3B, extensively used as autophagy biomarkers, to be significantly downregulated in miR-96-5p transfected cells compared to control mimic transfected cells. On the other hand, there was no significant difference in miR-96-5p inhibitor-treated compared to the control-treated cells. Our results clearly indicated miR-96-5p to be associated with autophagy protein LC3A/B. Therefore, upregulation of miR-96-5p in old muscle may partly be responsible for the decline in autophagy in aged skeletal muscle (Figure 10).

**Figure 10.**
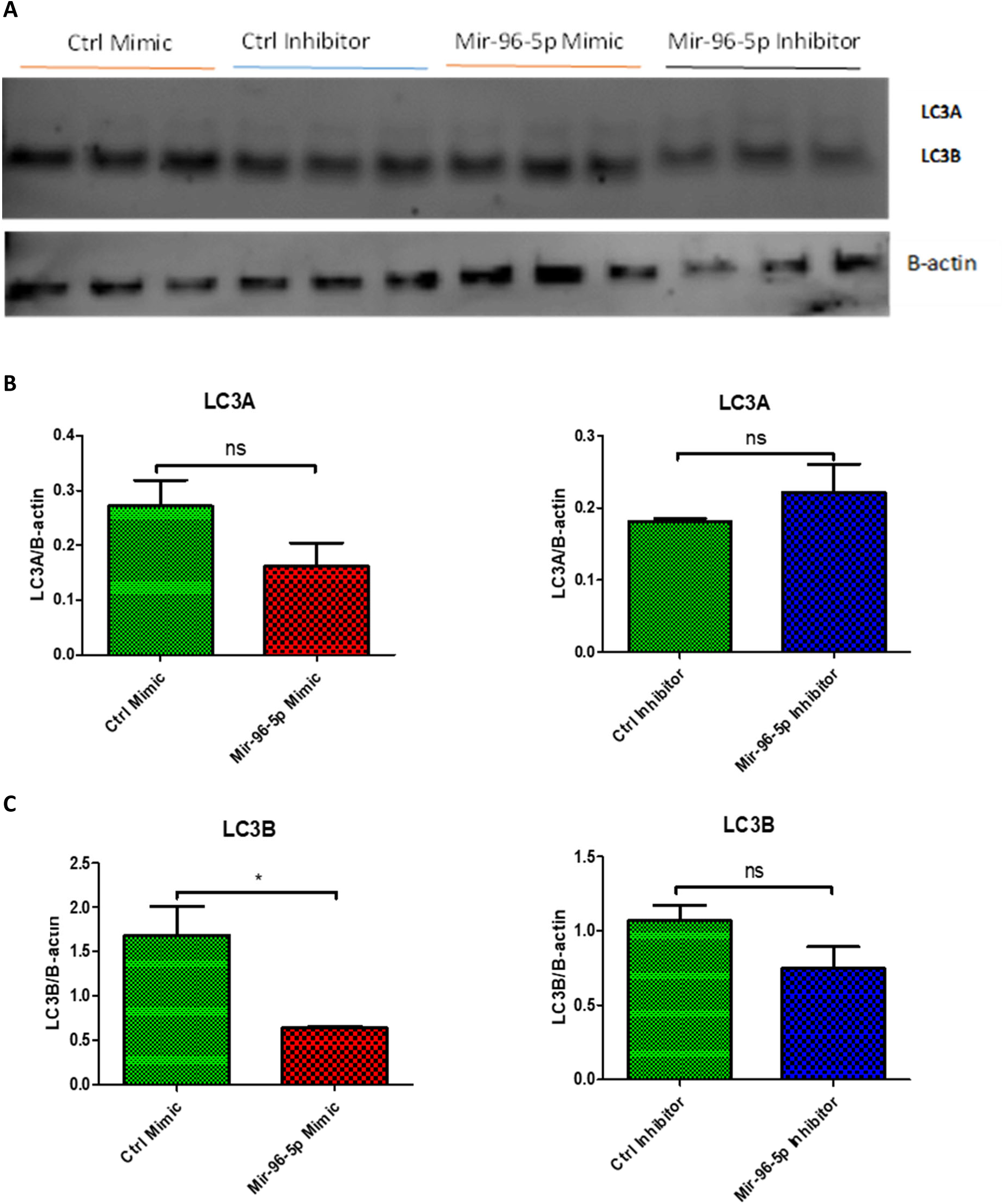
MiR-96-5p reduces autophagy markers in C2C12 myoblast cells. Cells were transfected with 50 nM of control mimic, miR-96-5p mimic, or 100nM control inhibitor or miR-96-5p inhibitor. Proteins were isolated 24h post transfection and western blot was conducted. **A.** showing western blot image against LC3A/B upon miR-96-5p treatment, and beta actin. **B.** showing quantification of western blot for LC3A. **C.** showing quantification of western blot for LC3B. Error bars represent mean ± SEM, n = 3, ns denotes not significant,*=p-value<0.01 to 0.05. In comparison to b-actin, the relative protein expressions were determined.

### Effect of miR-96-5p on OGDH expression and function

**Figure 11.**
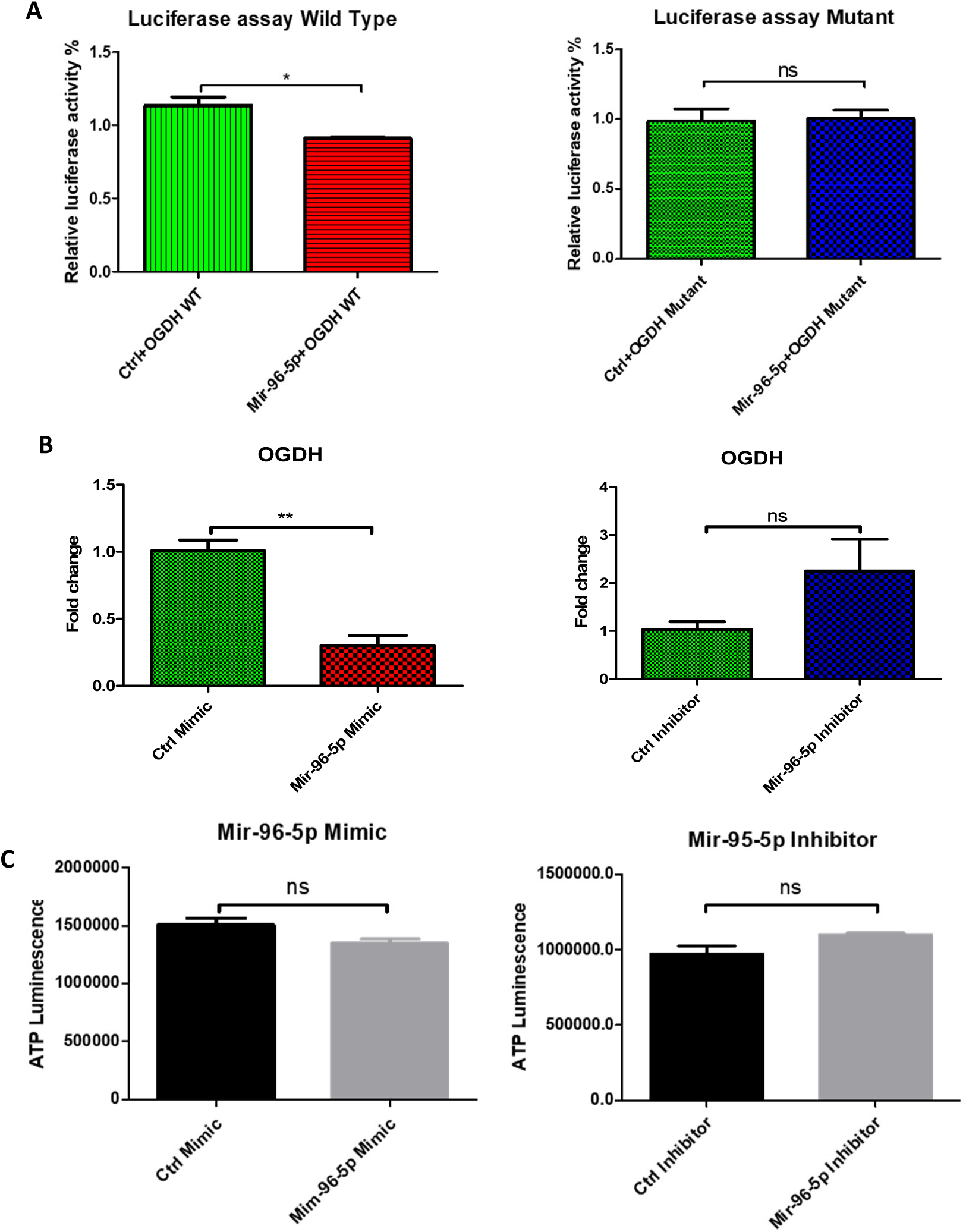
Interaction between miR-96-5p and OGDH. (A) Luciferase assay, using scRNA control or miR-96-5p co-transfected in pmirGLO vector with 3’UTR segment of OGDH of wild-type (WT) or Mutant nucleotides. Cells were analysed after 24h of transfection with 1μg of WT or mutant plasmid and 100nM of scRNA control or miR-96-5p mimic for luciferase activity using the Dual-Glo luciferase assay system and a luminometer. Fruit fly luciferase activity was normalised against renilla activity. (B) RT-qPCR analysis regarding the relative expression of OGDH for each treatment condition. (C) Mitochondrial ToxGlo assay was used to determine cellular ATP levels in cells treated with miR-96-5p mimic relative to the control mimic. Error bars represent mean ± SEM, n = 3, ns denotes not significant, *= p-value <0.01 to 0.05.

Quantitative PCR revealed cells transfected with miR-96-5p to have significantly lower expression of OGDH compared to cells transfected with the mock mimetic. According to qPCR, the miR-96-5p inhibitor appeared to have no effect on OGDH expression (Figure 20A).

Target scan using the https://mirdb.org/custom.html database (Wong and Wang, 2015) revealed significant alignment between the 3’UTR segment of OGDH and miR-96-5p in several species. This suggested a direct interaction to be responsible for the effects of miR-96-5p on OGDH expression. We sought to confirm this using the luciferase reporter assay, which revealed cells transfected with miR-96-5p to display reduced luciferase activity compared to cells transfected with the mock mimetic. This effect was only present in cells containing the wild-type OGDH segment but not those containing the mutant OGDH segment (Figure 20B).

Lastly, ATP assay rendered no significant effects between groups with regard to ATP production. Saying this, there was a non-significant trend towards lower ATP production in the miR-96-5p-transfected cells and higher ATP production in the miR-96-5p inhibitor-transfected cells, when compared to the mock mimetic and mock inhibitor transfected cells respectively (Figure 20C).

## Discussion

While the exact mechanisms involved in sarcopenia are complex and not fully understood, perturbed epigenetic and post-transcriptional regulation of gene expression likely plays an important role. In particular, dysregulated miRNA expression has previously been shown to be involved (Jung et al. 2017; Drummond et al. 2011). Furthermore, caloric restriction, a potent anti-ageing intervention, appears to ameliorate age-related muscular dysfunction, in part, through reducing age-related changes in the miRNA (Margolis et al. 2016; Mercken et al. 2013). Our study aimed to explore age-related miRNA expression changes in rat skeletal muscle, comparing young, old, and caloric restriction-treated old groups, to uncover how miRNA may influence muscle ageing and how caloric restriction could potentially slow down this process.

### Differential analysis of miRNA between young vs old and old CR

We identified 84 miRNAs to be differentially expressed in the gastrocnemius muscle of aged rats compared to young rats. We also found 109 miRNAs to be differentially expressed in old CR muscle compared to young rat muscle. This indicates CR to have an impact on the expression of miRNAs during the processes of muscle ageing. Compared to previous studies, several of the differentially expressed miRNAs we identified have been previously reported in muscle ageing RNA-seq studies. Our analysis partially replicated findings from previous human, mouse and rhesus monkey studies. We found 4 (miR-146a-5p, miR-483-5p, miR-675-5p and miR-539-5p) and 10 (miR-434-3p, miR-181c-5p and miR-411-5p, miR-483-3p, miR-146a-5p, miR-29b-3p, miR-369-5p, miR-434-3p,miR-136-5p and miR-411-5p) miRNAs to be differentially expressed both in our study and in studies of ageing in human (Soriano-Arroquia et al. 2016) and mouse (Jung et al. 2017; Kim et al. 2014) skeletal muscle. Furthermore, our results found one (miR-495) miRNA that has previously been shown to be differentially expressed in aged Rhesus monkey skeletal muscle (Mercken et al. 2013).

### GO and KEGG Pathway Analyses of miRNA Target Genes

GO analysis revealed miRNAs upregulated with age to downregulate genes involved in skeletal muscle tissue development, fatty acid oxidation regulation, and hexose metabolism, as well as cGMP-PKG AMPK, longevity regulation, insulin resistance, and pro-autophagic pathways. This suggests miRNA to be involved in age-associated impairments in muscle cell differentiation and metabolism. Furthermore, our result implicates miRNA overexpression in skeletal muscle in the downregulation of several longevity-associated processes related to insulin sensitivity, autophagy and nutrient sensing.

In addition, GO analysis revealed miRNAs downregulated with age to likely result in upregulation of genes involved in pro-inflammatory processes. Interestingly, our results suggested age-associated inflammation to result from the upregulation of complement and coagulation genes. The complement system is a proteolytic cascade that functions as a mediator of innate immunity, and the coagulation system plays a key role in haemostasis, serving as the body’s first line of defence against harmful stimuli and intruders. The activation of both result in inflammatory responses, therefore, maladaptive levels of activity can lead to aberrant inflammation (Esmon 2005). This may explain the link between upregulation of complement and coagulation in aged muscle.

Lastly, GO and KEGG pathway analysis was applied to previously undiscovered miRNAs differentially expressed with age. Results of these analyses suggested these novel miRNAs to target the PI3K-Akt, MAPK, mTOR and BMP signalling. Many of these signalling pathways have previously been shown to be important in optimal muscle function and the ageing process. For example, both MAPK and mTOR signalling have been shown to be important in determining satellite cell fate by promoting myogenic proliferation and differentiation (Keren et al. 2006; Segalés et al. 2015; Yoon 2017). Therefore, the miRNAs discovered in this study may contribute to the age-associated depletion of the satellite cell pool (Shefer et al. 2006) and could be investigated for therapeutic potential. Notably, however, the genes purportedly targeted by these novel miRNAs were predicted based upon sequence homology. Therefore, it is crucial that both the existence and targets of these miRNAs be validated in vitro.

### Impact of CR on the expression and function of differentially expressed miRNA in old skeletal muscle and their function

Our findings are in line with a previous publication, where it was demonstrated that CR was able to reverse the expression of age-associated altered miRNAs in rhesus monkeys (Mercken et al. 2013). Specifically, we found CR to normalise the expression of 45.5% of miRNA displaying dysregulated expression with age. Interestingly, our results show CR to have had a more profound impact on upregulated miRNAs. Indeed, GO and pathway analyses found the age-associated miRNAs, which CR suppressed towards youthful levels, to target many of the biological processes discussed earlier, such as those associated with skeletal muscle tissue development, metabolism and nutrient partitioning. Furthermore, through the suppression of miRNAs, CR seemed to upregulate longevity-associated AMPK and autophagy pathways (Ge et al. 2022). Notably, however, after FDR correction, the significance of the correlation between CR-suppressed miRNAs and these pathways was not maintained.

Turning to genes targeted by miRNAs rescued by CR, GO and KEGG pathway analyses found these to be largely associated with glucose metabolism. Reduced calorie intake is anticipated to cause a metabolic switch from carbohydrate to fatty acid oxidation (Bruss et al. 2010), therefore, this finding may be a mere artefact of this process. However, impaired glucose homeostasis (often leading to type 2 diabetes) is a well-characterised outcome of ageing (Kalyani & Egan 2013). Therefore, the miRNAs discovered to be upregulated by CR in this study may be important in understanding this symptom of ageing.

### Age-associated overexpression of MiR-96-5p negatively regulates differentiation, mitochondrial biogenesis, and autophagy

Based on bioinformatics and target analysis, we discovered that miR-96-5p is a key player in muscle ageing and a key mechanism in CR. This miRNA’s target genes were linked to biological processes such as insulin responses, positive regulation of fatty acid oxidation and skeletal muscle tissue development, as well as several longevity-associated pathways. Our results demonstrated miR-96-5p inhibition to significantly increase cell viability and markers of mitochondrial biogenesis (PGC-1α, COX I and TFAM) whilst miR-96-5p activity to significantly decrease markers of myogenic differentiation and autophagy, as well as TFAM expression. This was consistent with previous findings regarding the role of miR-96-5p in the negative regulation of myogenic differentiation (Nguyen et al. 2020). In addition, we demonstrated using the luciferase reporter assay, that miR-96-5p likely directly binds to the 3’UTR of OGDH and negatively regulates its expression. In addition, we postulated that overexpression of miR-96-5p would reduce ATP synthesis due to this suppression of OGDH expression. We, therefore, postulated that overexpression of miR-96-5p would reduce ATP synthesis due to suppression of OGDH expression. Although there was no significant difference, we observed there was a trend of reduced ATP production for cells treated with miR-96-5p mimic, and there was a trend of upregulation in ATP production in miR-96-5p inhibitor treatment group.

Even though our findings are intriguing, many questions remain unanswered, such as how miR-96-5p regulates mitochondrial function and biogenesis. Since OGDH silencing is causally linked to a decrease in mitochondrial biogenesis and function (Nemeria et al. 2014) and miR-96-5p has no obvious binding site on PGC-1, COX1 or TFAM mRNA, it could be hypothetically attributed to the repression of OGDH mRNA by miR-96-5p. Nonetheless, during CR, we discovered that miR-95-5p expression levels were equivalent to younger muscle expression levels, and OGDH was similarly recovered. As a result, we hypothesise that these positive effects of CR are caused by CR’s modulation of miRNA-96-5p in aged skeletal muscle so this should be further investigated.

## Conclusion

Our study revealed previously known and unidentified miRNAs to be differentially expressed in the muscle of aged rats, as well as which of these were responsive to caloric restriction (CR). Our results shed light on how miRNAs upregulated with age may contribute to satellite cell depletion, impaired myogenic differentiation, and metabolic dysfunction. These results were largely confirmed in our assessment of the effects of miR-96-5p in vitro. Furthermore, miRNAs downregulated with age were found to suppress a number of pro-longevity pathways, including AMPK and autophagy.

Interestingly, we also found CR to preferentially suppress miRNAs upregulated with age more so than rescue those downregulated with age. miRNAs suppressed by CR targeted many of the pathways targeted by miRNAs upregulated with age, including skeletal muscle tissue development, metabolism and nutrient partitioning, suggesting CR to upregulate these processes. However, CR was less able to downregulate the inflammatory-associated processes, thought to be dysregulated due to miRNA downregulation with age.

## Authors’ contributions

GA and JPM conceived the original idea. GA performed all the investigations and analysis. GA authored the original work, with CWB structuring the paper. The final version of the manuscript was edited and revised by GA, CWB, JPM, KGW and ASA. The study was supervised by JPM, KGW, and AV. PR and ASA assisted with both computational and laboratory investigation. BZ assisted with laboratory investigation.

## Conflicts of interest

JPM is CSO of YouthBio Therapeutics, an advisor/consultant for the Longevity Vision Fund, 199 Biotechnologies, and NOVOS, and the founder of Magellan Science Ltd, a company providing consulting services in longevity science.

GA is CEO of Fagus Antibody services, a company providing custom antibody services to biotech and academia.

